# Geometric analysis of postnatal maturational transformations in brain’s functional connectome

**DOI:** 10.64898/2026.11.13.766350

**Authors:** Vyacheslav R. Karolis, Joseph V. Hajnal, Saad Jbabdi, A. David Edwards, Grainne McAlonan, Tomoki Arichi

**Affiliations:** Research Department of Early Life Imaging, School of Biomedical Engineering and Imaging Sciences, King’s College London, London, UK; Oxford Centre for Integrative Neuroscience, Nuffield Department of Clinical Neurosciences, University of Oxford; Department of Forensic and Neurodevelopmental Sciences, Institute of Psychiatry, Psychology & Neuroscience, King’s College London, London, UK; MRC Centre for Neurodevelopmental Disorders, King’s College London, London, UK; Paediatric Neurosciences, Evelina London Children’s Hospital, Guy’s and St Thomas’ NHS Foundation Trust, London, UK

**Keywords:** multi-view Laplacian embedding, functional hierarchy, connectivity gradient, birth, functional development, neonates

## Abstract

Birth marks a profound shift in sensory experience, posing a question: does postnatal brain function represent simply a protracted continuation of prenatal developmental programmes or reflect system-level reconfiguration in response to the demands of the postnatal environment? To answer this, we first characterised functional hierarchies of the newborn human brain, separating sensorimotor from visual, unimodal from associative, and parietal from auditory systems. We then implemented a geometric framework to map global transformations that shape functional connectome hierarchies. We show that greater postmenstrual age at birth was associated with emergence of the frontoparietal and salience networks at the tip of the unimodal–associative hierarchical axis; in contrast, greater postnatal age was associated with increasing polarisation of primary perceptual and sensorimotor systems across functional hierarchies. Sex differences in the neonatal connectome recapitulated patterns associated with postmenstrual age at birth and postnatal age, suggesting more advanced maturation of functional architecture in females than in males. Together, the contrasting effects of prenatal and postnatal age support a model of postnatal reconfiguration of brain function, thereby indicating that birth could mark a transition between distinct modes of the functional organisation.

## Introduction

Birth marks a profound shift for the brain and body. Separated from the protective and nourishing *in utero* environment with its specific fetal blood circulation and dampened external stimulation, the human neonate at birth must immediately sustain autonomous physiological function while adapting to the full extent of gravity and unaltered sensory stimuli. How human brain function responds to this dramatic transition remains largely unexplored, posing a key question: does postnatal maturation of the brain’s functional organisation reflect a protracted continuation of a prenatal program or follow trajectories that extend beyond those established prenatally?

There are substantial methodological challenges to answering this question. Importantly, misalignment of developmental stages at birth between humans and exemplar animal models^1,2^ limits direct translation from animal studies, thus preventing precise insight into the perinatal factors that alter neuronal signalling and could drive system-level changes in human brain function^3,4^. In the meantime, a direct study of fetal-to-neonatal transitions in functional connectivity using fMRI with human neonates would be strongly confounded by differences in the imaging hardware, acquisition, and data processing required for studying an individual *in utero* or *ex utero*. Thus even if differences in brain function are seen before and after birth, a significant challenge remains in attempting to distinguish methodological effects from true changes in underlying neurobiology^5^.

Here we propose that, even if direct measurement of a change in the brain function over the birth divide remains problematic, the presence of a *deviation* from an unobserved prenatal trajectory can be indirectly inferred by analytical means using neonatal data alone, by leveraging inter-subject variability of functional connectivity in terms of postmenstrual age at birth and postnatal age at scan. Specifically, we hypothesise (Fig. 1a) that, if postnatal maturation is simply an extrapolation of prenatal functional connectivity maturational trajectories, then the effects of a greater postmenstrual age at birth and postnatal age at scan would be undistinguishable; that is, their sum - postmenstrual age at scan - would contain sufficient information about variance in postnatal maturational trajectory. Conversely, if postnatal development represents a deviation from a trajectory of the prenatal development, then the effects of postmenstrual age at birth and postnatal age at scan would contribute to variance in maturational trajectory as two separate factors.

**Figure 1.**
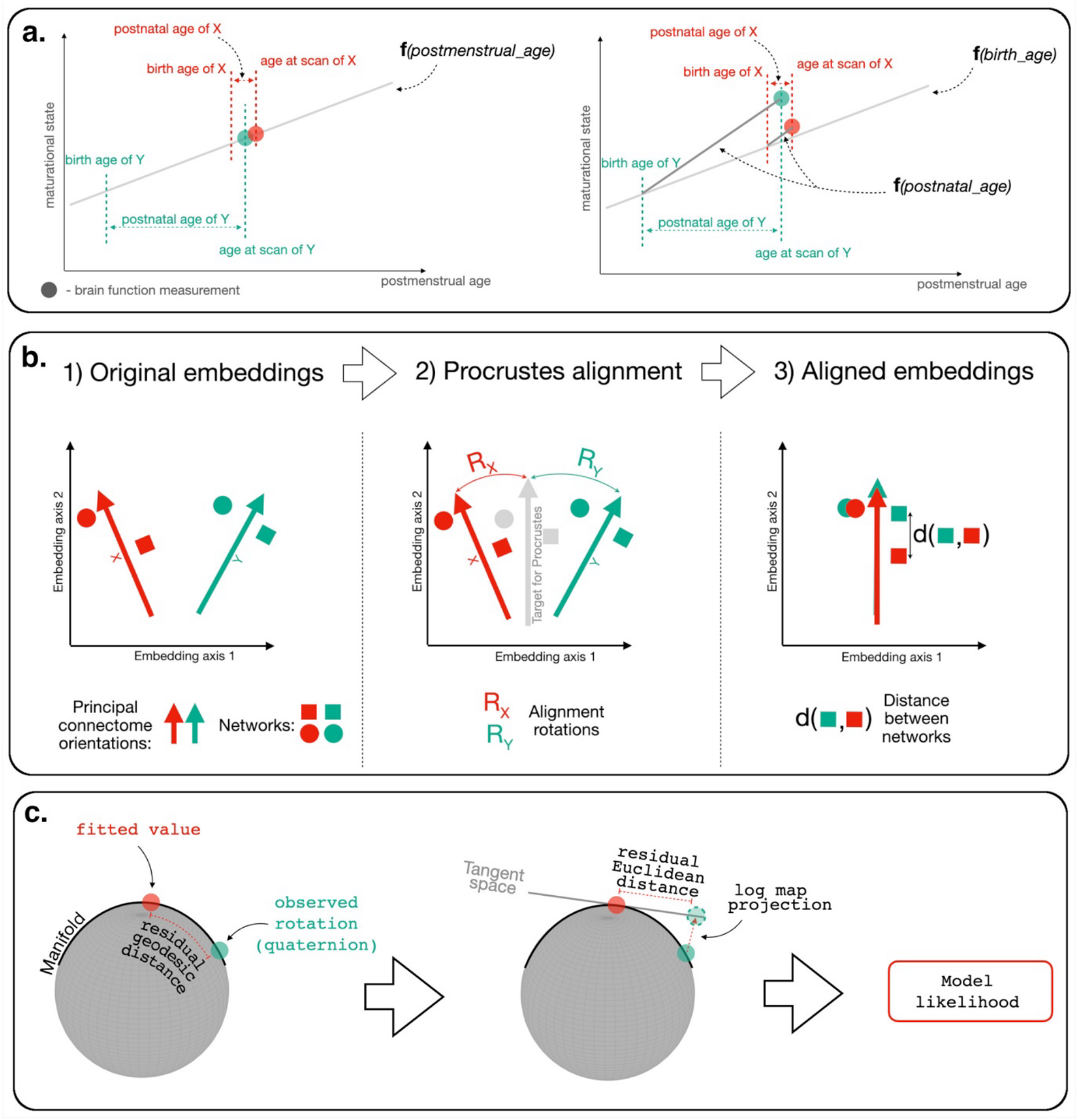
Methodological premises for the study. **a** Schematic representation of the predictions for two hypotheses on postnatal developmental trajectory. Bold “f” stands for “function”. X and Y are individual cases. If postnatal maturation represents a protracted continuation of the prenatal trajectory, the individual variability in the length of postmenstrual age at birth and postnatal age would not matter, as only their sum is important. Conversely, if postnatal maturation is not aligned with the prenatal trajectory, individual variations in postmenstrual age at birth and postnatal age would be associated with variations in the brain function that cannot be accounted by the sum of birth age and postnatal age. **b** The scope for the analyses of inter-individual differences. The multi-view embeddings allow modelling inter-individual differences in global connectome orientations, R_x_ and R_y_, whereas modelling inter-individual differences for the single-view embeddings are restricted to the analyses of the residual distances of network positions along embedding axes. X and Y are individual cases. **c** Modelling global connectome transformations. For this, rigid rotations from individual connectome embeddings to consensus embedding, calculated via Procrustes alignments, are converted into quaternions, i.e., 4D vectors. This permits modelling rotations as points on a 4D hypersphere with radius=1.

Early-life functional maturation is underpinned by rapid changes in structural connectivity ^6–9^, and thus is a multifaceted process^10^ whose characterisation would benefit from an integrated view of the evolving functional brain architecture. A common geometric formalism for summarising global relations in brain connectivity is to use Laplacian embedding of connectivity matrices, in which inter-regional connectedness is expressed as relative distance between pairs of brain regions in a low-dimensional Euclidean space^11^. Regions situated at the opposite poles of each embedding dimension define brain hierarchies, oftentimes called connectivity “gradients”^12,13^. This approach identifies the well-described segregation in the adult brain between unimodal sensory and transmodal association systems as a primary hierarchy, and between visual and sensorimotor systems as a secondary hierarchy^14^. In the neonatal brain^15,16^, several studies have shown that unimodal-to-transmodal hierarchy appears to be not yet present with the primary segregation axis instead separating sensorimotor and visual systems, which persists up to adolescent years ^17,18^.

Such findings however provide only a very coarse qualitative characterisation of the maturational process, limited to assessing the presence and ordering of “gradients”. An alternative - quantitative - approach has been to analyse maturational changes in the position of individual brain regions along embedding dimensions. However, in its current form, this approach loses a crucial aspect of the qualitative characterisation of maturation: global shifts in hierarchical organisation. By construction, each individual connectome is embedded in its own space, precluding direct comparison across individuals. As a result, prior to analyses of inter-individual differences, individual embeddings need to be aligned to a common target space. This implies that the global inter-individual shift in connectivity architecture, encoded in the magnitude of alignment transformation, is inherently lost and only its residuals can be analysed.

In contrast (Fig. 1b), the multi-view embedding approach for consensus-based characterisation of data structures enforces individual embeddings that span a shared sub-space^19^. Importantly, this enforcement is ambivalent to the global orientation of the individual embeddings, which can now become a legitimate target for analysis. In this study, we capitalised on this geometric property of multi-view embedding to bridge the gap, by linking the characterisation of hierarchical order with quantitative measures of intra-hierarchical organisation between the two previous approaches. We implemented a modelling approach that is compliant with their intrinsic manifold geometry of global shifts in the functional connectome (Fig. 1c) and applied it to a large sample of neonatal fMRI data^20^, with the aim of discriminating between two competing hypotheses about the relationship between pre- and postnatal functional brain development.

## Results

Here we analysed the functional MRI data of 517 term-born neonates from the open-access developing Human Connectome Project (dHCP) cohort^20^. For each neonate, the spatial maps of 40-component group-ICA factorisation reported in ^21^ were used as seeds to obtain network timecourses in native functional space, for which pairwise partial correlations were aggregated into connectivity matrices. For embedding, we retained a connectome of 29 networks (Fig. 2a), discarding networks that showed signatures consistent with a probable non-neuronal origin or contamination. The labelling of the networks in the figures below refers to the order number of the component in the entire 40-component set.

**Figure 2.**
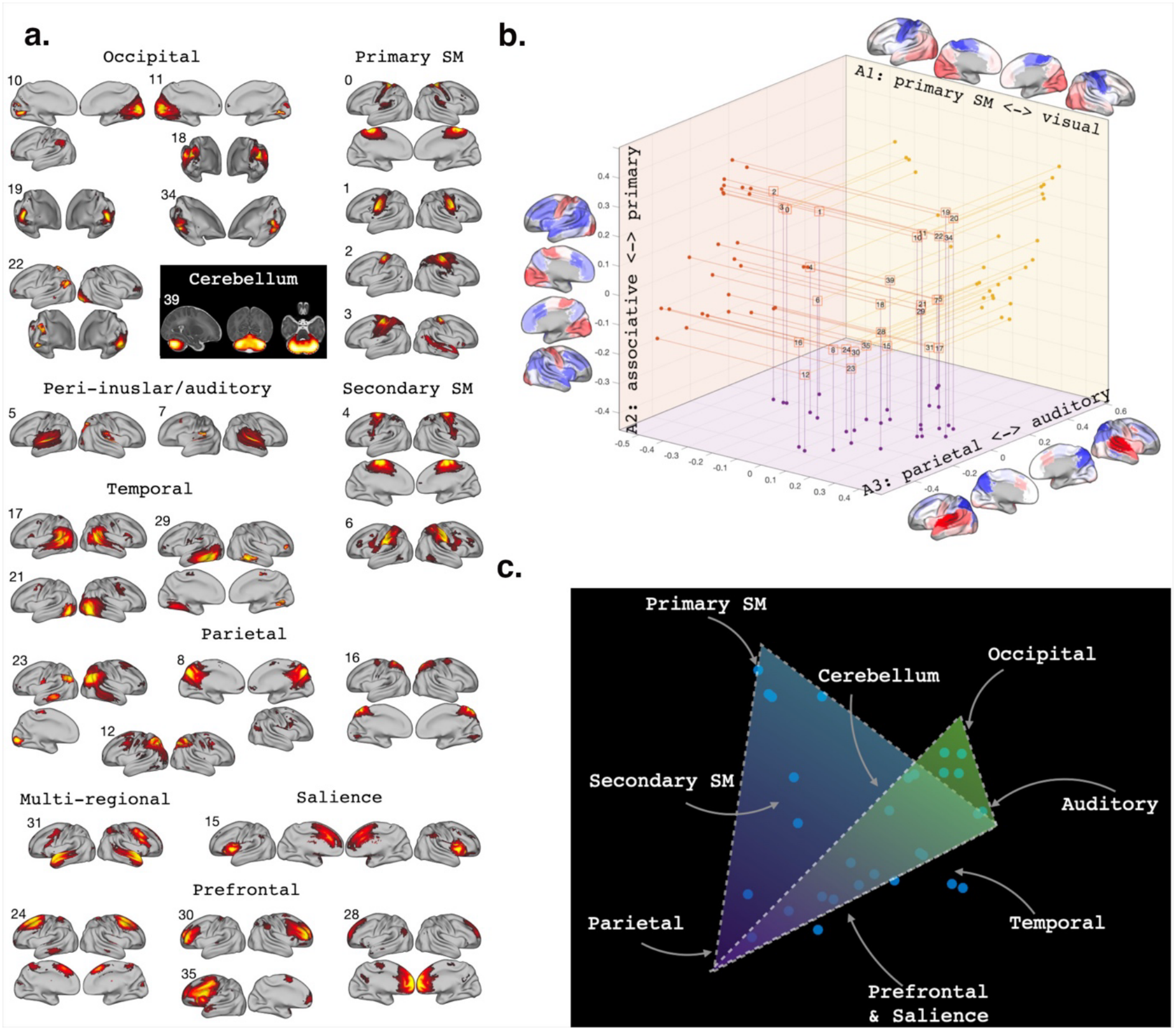
Neonatal functional networks and their consensus neonatal connectome embedding. **a** The spatial maps of resting-state neonatal networks used in the study. The maps were derived in the volumetric space; projections on cortical surface templated are used to aid visualisation only; **b** Consensus embedding and its projections onto 2D planes; the surface maps were produced by using networks to create winner-take-all parcellation, weighted by corresponding coordinates along a principal axis; **c** Schematic representation of embedding’s 3D geometry.

### Global characteristics of neonatal connectivity architecture

The consensus embedding of the neonatal functional connectome is shown in Fig. 2b-c. Its 3D representation reveals a meaningful anatomical architecture, with 4 distinctive poles in its structure: 1) primary sensorimotor (SM), 2) visual (occipital), 3) auditory (posterior peri-insular) and 4) parietal. We denominate the 3 principal axes of functional segregation, or hierarchies, in the neonatal brain as: 1) “primary SM <-> visual”; 2) “primary <-> associative” (i.e., primary SM+visual vs. prefrontal+parietal+temporal); 3) “parietal <-> auditory”. Other notable features in the embedded connectome include: an approximately equidistant location of cerebellum to all poles; the intermediate position of secondary SM networks (labels #4 and #6) between primary sensorimotor areas and associative networks; branching of the visual system into the two routes, ventral temporal (#21 and #29) towards the temporal/prefrontal cortices and superior occipital (#18) towards the parietal cortex.

### Modelling global maturational transformations

We characterised the global transformations in functional connectivity architecture by modelling rotations of individual connectomes to the consensus embedding. For this, the rotation matrices that encode rigid transformation between individual and consensus connectomes embeddings were converted into their quaternion representation, which we modelled as geodesic points on a 4D hypersphere with radius = 1.

The two-stage model comparison procedure (likelihood-ratio test for nested models, e.g., postmenstrual age at scan vs. postmenstrual age at birth & postnatal age, and Vuong tests for models with an equal number of predictors, e.g., postmenstrual age vs. postnatal age) was used to select the best performing model for global connectome reorganisation, whereby we: 1) defined a baseline model by comparing the models that contained sex and/or mode of delivery (natural birth vs. Caesarean) against the intercept model, so as to retain only the features in the baseline model that explained additional variance beyond the mean and 2) used this baseline model for systematic comparisons between models that contained age-related variables. To support the discontinuity hypothesis, the model that contained birth age and postnatal age at scan as separate variables had to outperform both the baseline model (incorporating appropriate Bonferroni corrections) and the model with postmenstrual age at scan as the variable. For completeness, the models that contained only birth age or only postnatal age at scan were also tested.

Both postmenstrual age at birth and postnatal age as separate variables accounted for the orientation of the connectome embeddings better than any other tested model (vs. baseline model, χ^2^ (6) = 26.22, Bonferroni-corrected p = .0008; vs. postmenstrual-age-at-scan model: χ^2^ (3) = 7.95, p = .047) (Fig. 3a). Importantly, the shared variance between birth age and postnatal age was 1.5%, indicating an independent contribution of each factor to model fitting.

**Figure 3.**
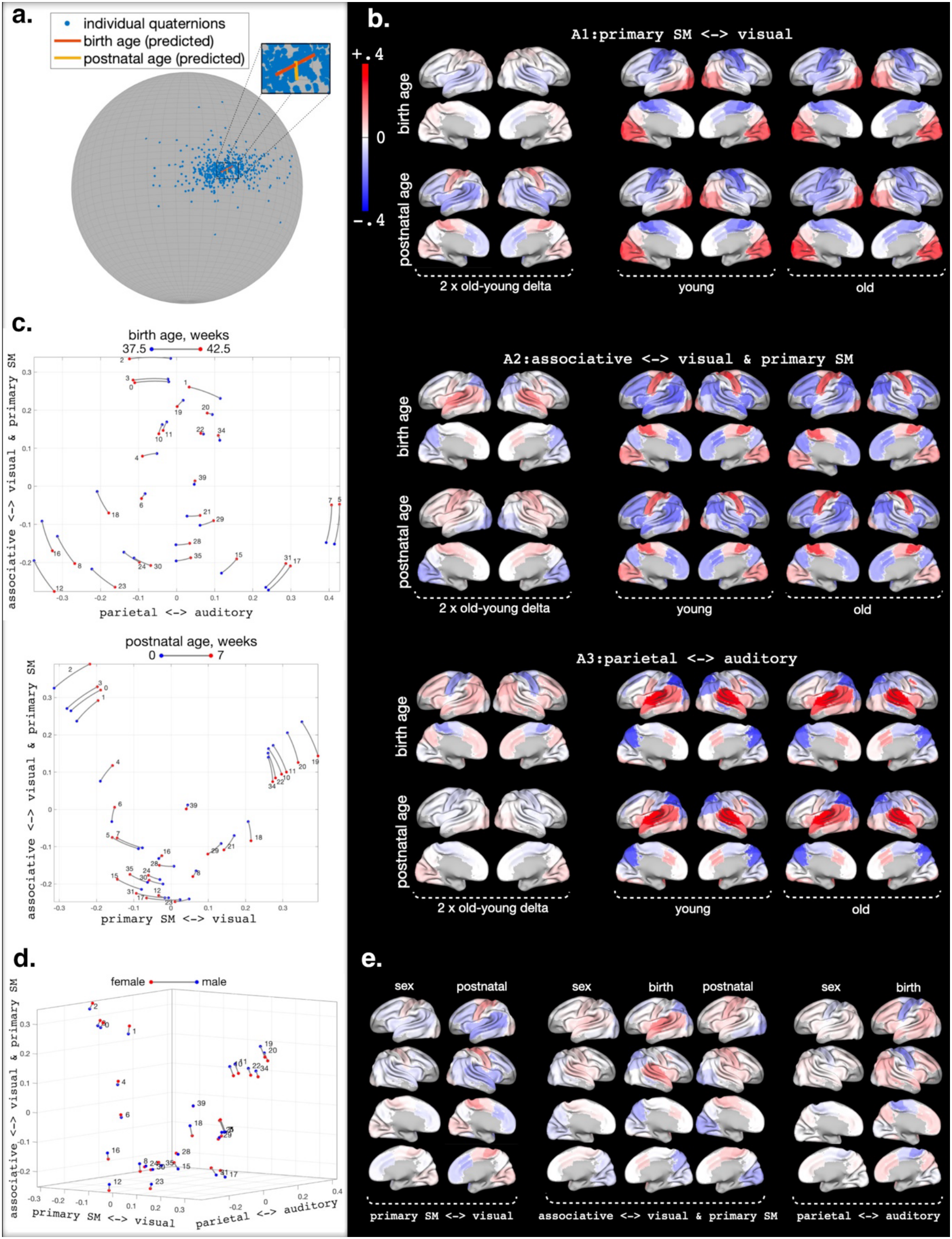
Global reorientation of connectomes in the embedded space. **a** Modelled maturation-dependent rotations (showing the first 3 quaternion values); (C) The effect of the modelled rotations on the network trajectories in the 3D embedded space; **b** spatial maps of one-dimensional projections of the maturation-dependent shifts (“delta”) along each principal axis (left – note that 2 x delta is shown to aid visualisation of the changing patterns) and network configurations for young and old ages (right). For birth age, young = 37.5 postmenstrual weeks and old = 42.5 postmenstrual weeks; for postnatal age, young = 0 postnatal weeks and old = 7 postnatal weeks. **c** Dominant in-plane rotations. **d** Effect of sex on global reorientation of connectomes in the embedded space. **e** spatial maps of one-dimensional projections of the 2 x “female-male” delta along each principal axis, presented alongside with the maturation-related shifts for visual comparison. The scale is the same as in b.

Fig 3b shows the spatial maps of the resulting shifts by birth age and postnatal age along each principal axis separately. These can be summarised as follows (for the per-network estimates of the shifts along each axis - see Supplementary Fig. 1):

1. ***“Primary SM <-> visual” axis*** The reshaping along this axis was dominated by *postnatal age*, resulting in transformation of this axis from polarisation of primary visual vs. SM networks towards a polarisation of primary visual networks vs. SM (both primary and secondary) and peri-insular (anterior insular salience and auditory) networks in individuals with older postnatal age.
2. ***“Associative <-> visual & primary SM” axis*** Both *postmenstrual age at birth* and *postnatal age* contributed to the reshaping along this axis but in different ways. For greater *postmenstrual age at birth*, this resulted in reshaping of the axis towards a polarisation of externally oriented (frontoparietal and salience) networks vs. primary visual & SM networks, with auditory networks and adjacent superior temporal networks shifting towards a more intermediate location along the hierarchy. For older *postnatal age*, we observe a differentiation between occipital and primary SM networks, resulting in transformation of this axis towards a polarisation between associative and primary SM networks, with primary visual networks re-mapping towards a more intermediate position.
3. ***“Parietal <-> auditory” axis*** Restructuring along this axis was dominated by *postmenstrual age at birth*, preserving the polarisation between parietal and auditory networks but resulting in a remapping within the existing hierarchy, with a convergence of associative networks towards the auditory pole and convergence of primary SM towards parietal pole in individuals with greater postmenstrual age at birth.

Alternatively, given that the shifts due to the effects of birth age and postnatal age were predominantly confined to remapping along two axes each, the dominant patterns of re-alignment associated with each variable can be meaningfully demonstrated as network trajectories in 2D planes, as shown in Fig. 3C. This representation of the result is particularly useful for the effect of postnatal age, illustrating that its increase leads to the re-alignment of “primary SM <-> visual” and “associative <-> visual & primary SM” axes towards a 3-way polarisation, whereby the associative networks no longer occupy an intermediate position between primary SM and visual networks, but rather form mutually orthogonal relations to them.

### Effect of sex on embedding organisation

The baseline model for the analysis of maturational factors on global re-organisation included sex (vs. intercept model: χ^2^ (3) = 15.70, Bonferroni corrected p = .0039). Fig 3d shows the effect of sex on network alignment in the 3D embedded space. Notably, females tended to demonstrate features of a more mature functional connectome, as the change from male to female connectome architectures aligns with transformations associated with greater birth and postnatal ages (Fig 3e). In particular, we observe: a) along the “primary SM <-> visual” axis, a greater convergence of the associative and auditory networks towards the primary SM pole; b) along the “associative <-> visual & primary SM” axis, a divergence between occipital and primary SM networks, with occipital networks converging towards the opposite, associative, pole; c) along the “parietal <-> auditory” axis, a convergence of primary SM networks towards the parietal pole.

### Modelling individual network trajectories

For completeness, to model the residual transformations that are not captured by the global transformation analysis, we also analysed our data using the standard “connectivity gradient” approach ^12^, whereby the single-view-derived individual embeddings are aligned to the consensus-based embedding prior to analysis of individual network trajectories within the embedded space. Postmenstrual age at scan was the best predictor for the individual network trajectories in 10 out of 14 cases where any model that contained one of the age-related factors outperformed the baseline model (Fig 4). The remaining 4 models contained both birth age and postnatal age as predictors. Whereas a comprehensive summary is complicated due to a diversity of the individual network trajectories, we note several features: a) a further divergence of auditory networks (#5 & 7) and primary visual networks (#10, 11 & 19) towards the corresponding ends along, respectively, the “parietal <-> auditory” and “associative <-> visual & primary SM” axes; b) a shift of the cerebellum (#39) to a central position in the embedded space; 3) the lack of age-related significant effects in the proximity of the parietal pole.

**Figure 4.**
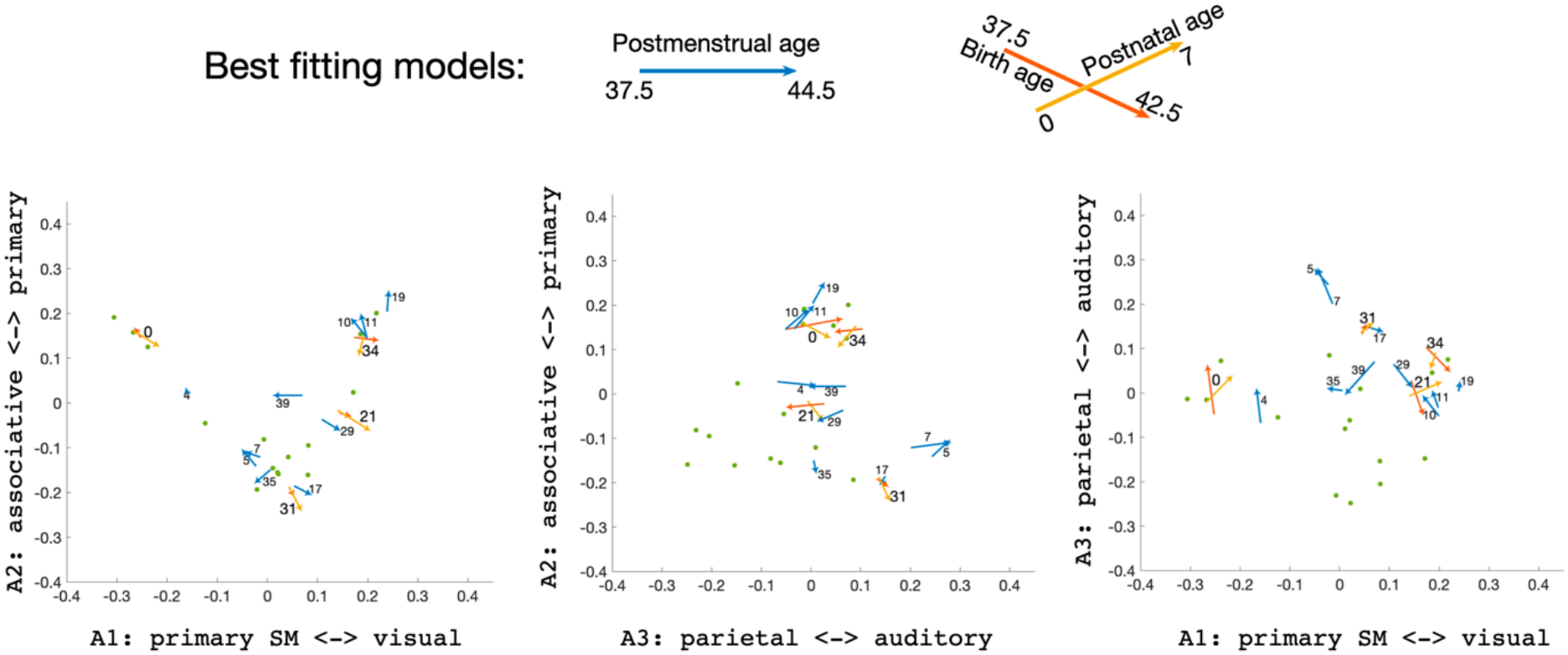
Projections of Individual network trajectories to the 2D plane space. For networks, which trajectory was better accounted for by birth age and postnatal age, the contribution of each variable is shown separately. The networks for which neither model passed a significance criterion against the baseline are shown as dots.

## Discussion

In this study, we characterised the large-scale functional architecture of the neonatal brain and its postnatal transformation using a bridging framework between two approaches for the description of maturational changes in brain hierarchies: the qualitative global, defining the existence and ordering of the hierarchies, and local quantitative, capturing residuals of the global transformations in positioning and/or expressiveness of networks along predefined functional hierarchies. In contrast, our framework enables quantitative characterisation of the maturation-related global transformations in the brain hierarchies by modelling rotations of the principal embedding axis of the functional connectome. We first described a sample-average representation of the neonatal brain’s hierarchies, identifying three axes of functional segregation in the neonatal brain: (i) “primary SM <-> visual”, (ii) “primary perceptual <-> associative”, and (iii) “parietal <-> auditory”, with four distinct poles: primary SM, occipital, auditory, and parietal. We then performed the analysis aimed at answering the study’s key question of whether postnatal functional development simply reflects a continuation of the protracted prenatal developmental programme. Importantly, we found that both postmenstrual age at birth and postnatal age contribute independently to maturational trajectory in hierarchical transformations, explaining variance beyond those accounted for only by their sum. These findings are thus in line with the hypothesis that functional re-organisation following birth is not just a uniform continuation of prenatal maturational patterns. Specifically, greater postmenstrual age at birth was associated with the emergence of the frontoparietal and salience networks at the tip of the unimodal-associative hierarchy; whilst older postnatal age led to redefining of the hierarchical poles along visual-sensorimotor and unimodal-associative hierarchies, towards hierarchies with visual separating from both associative and sensorimotor systems and sensorimotor from associative systems. Finally, we found that differences between the female and male connectome recapitulate motifs associated with greater postmenstrual age at birth and postnatal ages, which suggest that on average, the female neonatal functional connectome architecture tends to demonstrate more advanced maturation compared to its male equivalent. No significant effect of mode of delivery was observed.

Beyond characterisation of the networks that define the poles of the main axes of segregation, our analysis also revealed meaningful anatomical relations between hierarchical poles and intermediate networks along the hierarchy, showing a hierarchical transition from primary SM to higher associative regions via secondary SM and a segregation of visual information flow into the canonical dorsal and ventral visual streams via the superior occipital and ventral temporal networks. We also note a special role of cerebellum, which is uniquely positioned with the shortest mean distance in the global embedding space, aligning with both contemporary views that it is broadly connected enabling support of a wide spectrum of functions, including motor coordination, cognition, and emotional regulation^22,23^, and that its structural connections emerge very early in development^24,25^.

Useful comparisons of our results, highlighting both similarities and discrepancies, can be made with the results derived in previous studies using the “connectivity gradient” framework^10,14,16,20^. Firstly, corroborating the results of an earlier study derived using the first release of the dHCP data of 40 neonates^15,16^, we observe segregation between visual and sensorimotor areas within a primary functional segregation in the neonatal brain, which in the adult brain is typically reproduced as the second gradient axis. This finding is also in line with studies by Dong et al.^18^ and Nguyen et al.^17^, which suggest that a switch of principal axes ordering to the canonical adult model does not occur before the onset of adolescence. Second, we observe functional segregation at the opposite ends anchored at primary networks and associative networks., reminiscent of the unimodal-to-transmodal^11^ (alternatively, known as sensory-association^27^) primary axis in the adult brain “connectivity gradient” literature. In the adult brain, the transmodal pole is anchored at the default mode network, bordered by the frontoparietal networks, and the unimodal pole is anchored at the sensorimotor and visual networks, bordered by auditory networks^27^. In marked contrast, in neonates we find little segregation between parietal, temporal, and prefrontal networks along this primary-associative axis and the posterior peri-insular networks (overlapping with the auditory cortex) which are positioned at the associative pole of this brain hierarchy. We found that early segregation amongst associative networks occurs along the third, parietal-auditory, axis, with prefrontal and salience networks bordering the parietal pole and temporal networks bordering the auditory pole. Overall, in comparison to the adult connectome, the most prominent features of the neonatal functional connectome are a segregation between visual, auditory and sensorimotor systems and the lack of a coherent default mode system that would eventually crown the canonical unimodal-associative hierarchy that has been well described later in life.

Critically, the analytical approach that we developed allowed us to examine how the principal axes of functional organisation transform during the postnatal period and, with this, the network systems defining these principal axes. We show that the prenatal developmental trajectory, indirectly indexed by the transformations associated with increasing postmenstrual age at birth, progressively define a hierarchical axis spanning primary sensorimotor and visual systems to frontoparietal and salience systems, establishing the organisational polarity across externally oriented brain functions. In contrast to that, with increasing postnatal age, primary visual and sensorimotor systems become increasingly differentiated as distinct organisational poles, consistent with the idea of emerging specialised functional hierarchies that could support the demands of postnatal sensory and motor experience. Thus, the sequence of developmental events can be conceptualised as follows: prenatal functional development establishes an organisational scaffold for the interaction with the external environment and, with birth introducing the brain to the ex-utero environment, postnatal development reorganises and accentuates this architecture by increasingly polarising primary sensorimotor and visual systems within their respective hierarchies.

Our results also provide novel insight into the potential developmental processes that may give rise to the unimodal-transmodal hierarchy of the adult brain, whereby the DMN anchors its transmodal pole. Our analyses show that regions constituting the hubs of the adult DMN are already embedded within the frontoparietal system in the neonatal period. This organisation is consistent with emerging evidence from fetal fMRI studies^5^ showing that putative DMN regions become progressively integrated into the frontoparietal system during late gestation. One possibility is that this integration represents an intermediate stage in DMN development, followed by progressive differentiation of DMN regions from the frontoparietal system as connectivity among them strengthens. In this regard, the frontoparietal system may provide an early organisational context from which the DMN subsequently differentiates into a functionally distinct system.

A striking result was that another biologically meaningful variable, sex, was associated with transformations that suggest advanced maturation of the female connectome around perinatal period. Specifically, the effect of sex was in alignment to the effect of postnatal age along the primary SM-visual axis and the effect of postmenstrual age at birth along the parietal-auditory axes, with a mixture of the two along the unimodal-association axis, with female connectomes showing features associated with older postnatal age along each axis. Although sex differences in brain maturation have been reported across later developmental stages, our findings show that patterns characteristic of more advanced functional hierarchical organisation in females are already evident in the neonatal period.

Finally, using the standard approach^12^, modelling the residual displacements of individual networks within the connectome, we found that these displacements were more often better accounted for by the postmenstrual age at scan alone. Summarising the local refinements is complicated by their diverse presentation. However, qualitative assessment suggests that certain patterns are in line with those found in the analyses of global transformations; specifically, we observe a trend towards the segregation of primary sensory networks. We also note a particular case where analysis of the relative trajectories of individual networks is more informative than the analysis of the global transformations. Specifically, because of its central location, global transformations have little effect on the assignment of the cerebellum along any axis. In the meantime, the analysis of relative trajectories shows even closer alignment of the cerebellum to the centre of the embedding space, thereby characterising its role as a truly multi-functional component in the neonatal connectome.

The above results indicate that local and global aspects of neonatal connectome reorganisation may be differentially associated with maturational factors. Thus, functional development may proceed through both refinement within established hierarchical organisation and reconfiguration of the organisational axes themselves, with the former predominantly reflecting a continued progression of the prenatal maturational trajectory of developmental organisation. An alternative interpretation is however possible based on statistical considerations. Whereas the composition of the current dataset was near-optimal for estimating the independent contributions of the postmenstrual age at birth and postnatal age factors (only 1.5% shared variance), it is intrinsically disadvantaged for comparison against the restricted model, in which the sole predictor, postmenstrual age at scan, is the sum of the two factors in the full model. The additional explanatory information available to the two-factor model is therefore necessarily limited to variation in postmenstrual and postnatal age that is not captured by postmenstrual age at scan, which itself necessarily shares substantial variance with each of the two component factors (39.4% and 48.3%, respectively). Consequently, the differential effects of the two maturational factors need to be sufficiently strong to overcome the bias towards restricted models.

A limitation of our analysis is that the predictions are built in the absence of the ground truth about the trajectory of prenatal development. Here its extrapolation into the neonatal period is approximated as a function of postmenstrual age at birth which, of course, does not account for the potential changes in brain function that may occur over the actual fetal-to-neonatal transition, an unknown “birth delta”. However, the absence of this ground truth is tangential to the core of our conclusions. This instead leaves an open question about the main contributing factors (birth transition vs postnatal adaptation), their interaction, and about the magnitude of the shift compared to their prenatal status. Crucially, the observed differential effects of postmenstrual age at birth and postnatal age on the trajectory of functional development points to a major limitation of studying preterm infants to accurately mirror prenatal development. The definitive answer to this open question thus needs to be achieved through future studies involving robust harmonisation of fetal and neonatal observations. This would require accounting comprehensively – and non-trivially - for a concurrent and profound shift in the observational mode that also occurs with birth, due to the enormous inherent practical and methodological differences of imaging in the womb versus acquiring images directly from a sleeping neonate.

## Methods

### Participants

Participants were recruited as a part of the developing Human Connectome Project^28^ which was approved by the National Research Ethics Committee (14/LO/1169) and informed written consent given by the parents of all participants. The sample was composed of the entire set of term-born neonates in the dHCP cohort (517 babies, 54% male, postmenstrual age at birth > 37 weeks). The age at scan ranged from 37.4 to 44.9 postmenstrual weeks (Supplementary Fig. 2).

### Data and data analyses

The study utilises the open-access functional MRI neonatal release 3 data, released using the dHCP resting-state functional MRI (rfMRI) pre-processing pipeline for neonates^20^. In brief, each dataset consisted of 2300 volumes (45 slices each), acquired using single-shot EPI (TE/TR = 38/392, multiband factor = 9, flip angle = 34 ^°^), with slice matrix = 67 x 67 and 2.15 mm isotropic resolution. Pre-processing steps included correction for local field inhomogeneity using *topup*^29^, slice-to-volume motion correction and dynamic susceptibility distortio correction using *eddy*^30^, residual temporal artefact regression using FSL FIX^31^. The data were registered to the template space corresponding to 40 weeks of gestation using the dHCP registration pipeline, with warps between native functional and template spaces provided as part of the data release.

Group-level maps of functional networks were acquired using 40-component group-ICA^32^ from a partially overlapping sample of 311 neonates with mean postmenstrual age = 37.4 weeks^21^. The weighted-average network timecourses were computed in native fMRI space using thresholded (Z > 3) and normalised network maps as seeds. All analysis were performed in the volumetric space; the volume-to-surface mapping of the cortical network maps and spatial maps obtained in downstream analyses was used to aid visualisation.

Individual functional connectome matrices were formed using L2-regularised partial correlations, computed for the entire set of 40 maps using the FSLnets toolbox (https://web.mit.edu/fsl_v5.0.10/fsl/doc/wiki/FSLNets.html) with the default regularisation parameter (rho = .1). This was subsequently pruned to retain entries for 29 networks, with the remaining 11 maps discarded as showing signatures of a non-neuronal origin. Among 11 discarded maps, 5 maps were located in the ventral orbitorfrontal (vOF) cortex. Although these maps may partially reflect biologically meaningful signals, their striped spatial appearance, together with the known susceptibility of vOF regions to spatial distortions in gradient EPI sequences, justified their exclusion from further analyses.

The multi-view embeddings of the connectome were obtained as described in Kumar et al.^19^ (see Data availability statement for the location of the Matlab scripts) The regularisation parameter towards shared-space embedding was set such that it contained 99% of the mean data variance of the individual embeddings. We settle on the 3D embedding (after discarding the null dimension of embedding) because: 1) it allows for geometric analysis supported by visual assessment; 2) The 4D embedding preserved the characteristics of the first three dimensions, while separation along the fourth dimension reflected a segregation of the salience and adjacent ventral prefrontal cortex from functionally diverse anatomical systems (Supplementary Fig. 3). This implies that higher dimensions capture decoupling of finer-scale functional systems rather than additional axes of global functional organisation. Of note, the inclusion of the discarded vOF maps into the embedding resulted in them being a separate pole in the 4th dimension of embedding (Supplementary Fig, 4), while preserving the global embedding structure of the other regions in 3D. This peripheral role rendered further justification for their exclusion from the analyses.

The effect of maturational factors on the global shift of connectome orientation was assessed by analysing the rotation of the individual embeddings to the consensus embedding. For this, rotations were converted to their quaternion representations and modelled as geodesic points on a 4D hypersphere with radius = 1. The log map projections of residuals to the tangent hyperplanes at fitted values were used to compute Euclidian-metric likelihoods for the multivariate Gaussian with mean=0 and empirically estimated covariance matrix.

The two-staged model comparison procedure utilised likelihood-ratio and Vuong tests, respectively, for nested models (postmenstrual age at scan vs. birth age+postnatal age) and for models with an equal number of predictors (e.g., postmenstrual age vs. postnatal age). At the first stage, the effects of sex and mode of delivery (natural birth vs. pre-labour Caesarean) were assessed with respect to the intercept model (Bonferroni-corrected). The statistically best performing model was used as a baseline model for the second stage where explanatory power of birth age and/or postnatal age was assessed against that of their sum (i.e., age at scan). The birth-age and/or postnatal-age model was considered as best fitting if it outperformed both the baseline model (also Bonferroni-corrected) and the age-at-scan model.

Finally, using the standard single-view approach, we analysed the local age-related changes in connectivity, characterised by relative maturation-related displacements of networks within the embedding space, using multivariate regression of network x, y, z coordinates, following their Procrustes transformation to multi-view consensus embedding.

## Data availability

The data and code presented in this paper are available in the following locations:

1. dHCP fetal release: https://nda.nih.gov/edit_collection.html?id=3955
2. The network timecourses, the list of participants and Matlab code implementing multi-view embedding and geodesic fitting: https://gin.g-node.org/slavakarolis/neonatal_connectome_geometry
3. The neonatal spatio-temporal dHCP atlas: https://git.fmrib.ox.ac.uk/seanf/dhcp-resources/-/blob/master/docs/dhcp-augmented-volumetric-atlas-extended.md?ref_type=heads

## Author contributions

VK, JH, SJ, TA – Conceptualisation

VK – Methodology, Software, Formal Analysis, Visualisation, Writing-Original Draft

JH, DE, GM, TA - Writing-Review & Editing

GM, TA - Funding acquisition

## Acknowledgements

The data used in this study were acquired with the funding by European Research Council under the European Union’s Seventh Framework Programme (FP7/20072013)/ERC grant agreement no. 319456 (dHCP project). The results leading to this publication have received funding from the Innovative Medicines Initiative 2 Joint Undertaking under grant agreement No 777394 for the project AIMS-2-TRIALS. This Joint Undertaking receives support from the European Union’s Horizon 2020 research and innovation programme and EFPIA and AUTISM SPEAKS, Autistica, SFARI. GMM is part-funded by the National Institute for Health and Care Research (NIHR) Maudsley Biomedical Research Centre (BRC). TA was funded by a MRC Senior Clinical Fellowship [MR/Y009665/1] and receives support from the MRC Centre for Neurodevelopmental Disorders, King’s College London [MR/N026063/1]. The views expressed are those of the authors and not necessarily those of the any above-mentioned funder, the NIHR or the Department of Health and Social Care.

