## Supplementary Figures for "Geometric analysis of postnatal maturational transformations in brain’s functional connectome"

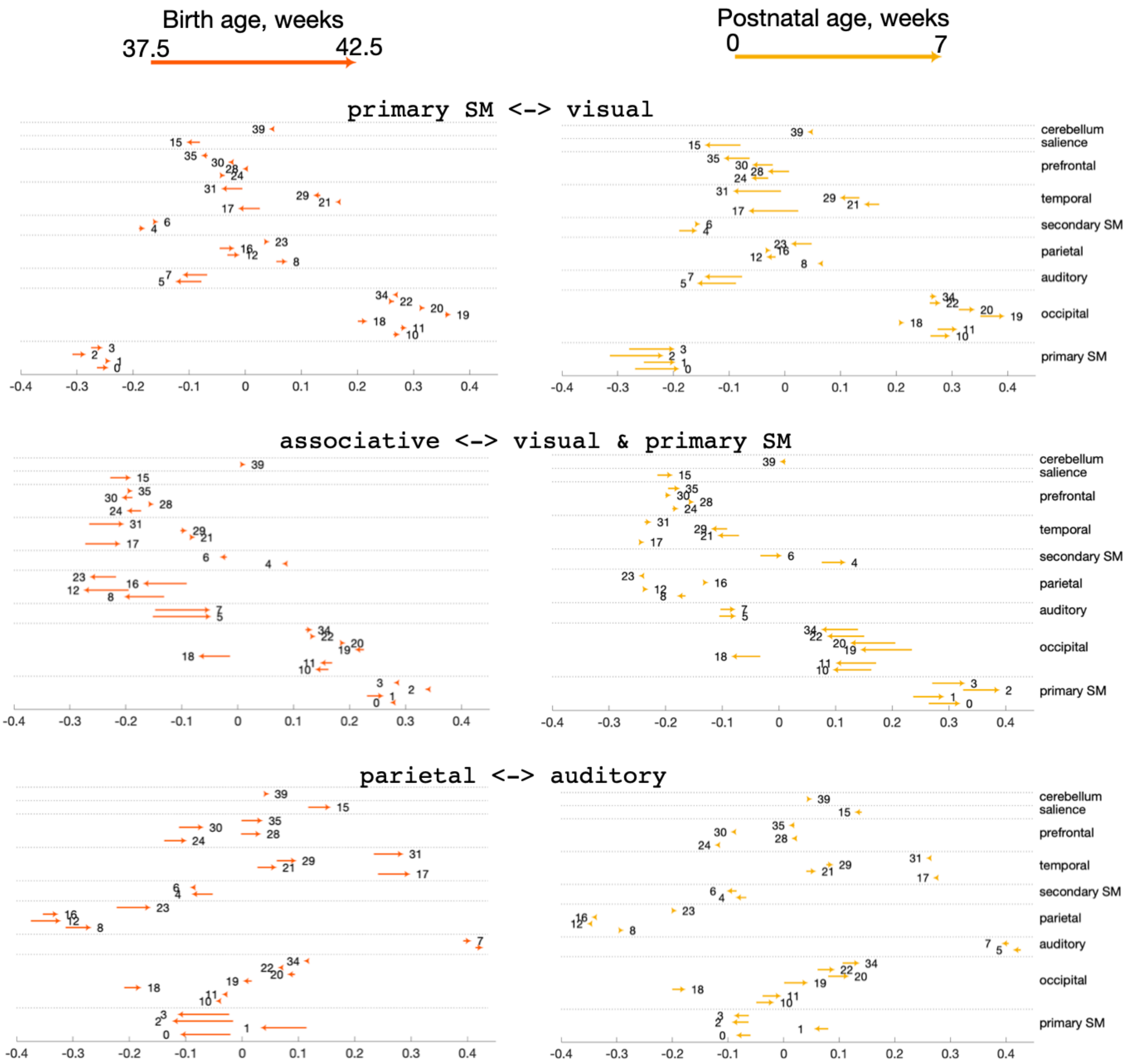


*Supplementary Fig. 1.* One-dimensional projections of the maturation-dependent shifts along each principal axis. Position of an arrow tail corresponds to the estimated network coordinate at a young age (0 weeks old for postnatal age and 37.5 postmenstrual weeks for birth age); position of an arrow tip corresponds to the estimated network coordinate at an older age (7 weeks old for postnatal age and 42.5 postmenstrual weeks for birth age).

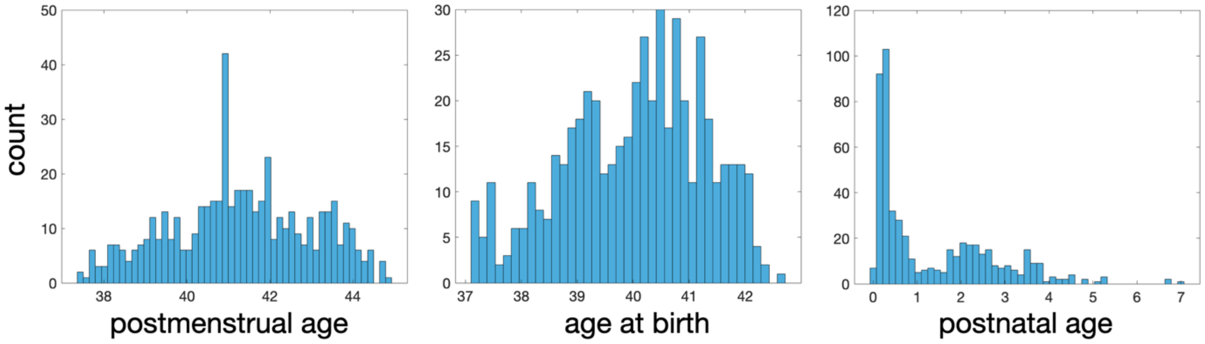


*Supplementary Fig. 2. Distribution of ages in the study sample.*

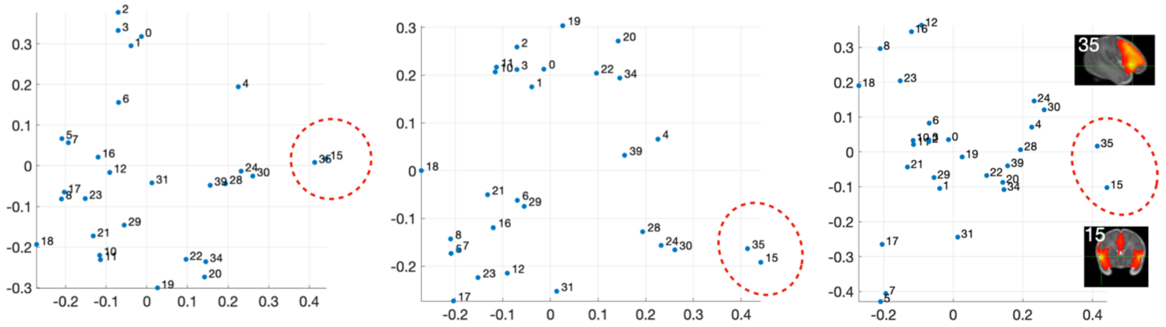

*Supplementary Fig. 3. Plane-projections of the 4th dimension of embedding (shown along x- axis). Salience and adjacent ventral pre-frontal networks are highlighted.*

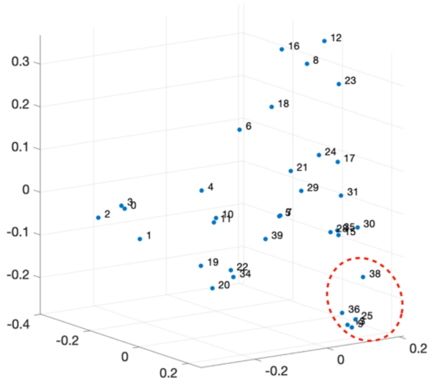

*Supplementary Fig. 4. Embedding with inclusion of ventral orbito-frontal networks*
